# SRSF3 knockdown-induced cellular senescence as a possible therapeutic strategy for non-small cell lung cancer

**DOI:** 10.1101/2025.05.05.652234

**Authors:** Shinji Nakamichi, Natalia von Muhlinen, Leo Yamada, Jilian R. Melamed, Tyler E. Papp, Hamideh Parhiz, Drew Weissman, Izumi Horikawa, Curtis C. Harris

**Affiliations:** Laboratory of Human Carcinogenesis, Center for Cancer Research, National Cancer Institute, National Institutes of Health, Bethesda, MD 20892, USA; Department of Medicine, Perelman School of Medicine at the University of Pennsylvania, Philadelphia, PA, USA

**Author notes:** Corresponding authors: Laboratory of Human Carcinogenesis, Center for Cancer Research, National Cancer Institute, National Institutes of Health, 37 Convent Dr., Bldg. 37 Room 3068A, Bethesda, MD 20892, USA.

**Keywords:** NSCLC, SRSF3, siRNA knockdown, cellular senescence, therapeutic strategy

## Abstract

Tyrosine kinase (TK) inhibitors improve clinical outcomes in non-small cell lung cancer (NSCLC) with targetable mutations. However, such NSCLC cases only consist of about 50% in the western populations. This study, for the first time in NSCLC cells including those without a targetable TK mutation, explores a tumor-suppressive activity of siRNA knockdown of a splicing factor SRSF3, which was reportedly effective in other cancer cell types. The knockdown of SRSF3 increased cellular senescence, indicated by senescence-associated β-galactosidase activity and reduced cell proliferation, in all NSCLC cell lines examined, including A549 (no TK mutation; *TP53* wild-type), NCI-H1975 (*EGFR* L858R/T790M; *TP53* R273H mutant), NCI-H322 (no TK mutation; *TP53* R248L mutant) and NCI-H596 (no TK mutation; *TP53* G245C mutant). An increase in apoptotic cleavage of caspase-3 and poly(ADP-ribose) polymerase was also observed in A549 cells. p53β, a tumor-suppressive p53 isoform generated via alternative mRNA splicing, was upregulated by SRSF3 knockdown, as previously reported in normal fibroblasts. However, neither cellular senescence nor apoptosis was increased by overexpression of p53β, suggesting no or minimum contribution of this p53 isoform to the tumor-suppressive activity of SRSF3 knockdown in NSCLC cells. Our gene expression assay indicated that the SRSF3 knockdown-induced senescence in NSCLC cells may be mediated by downregulation of TOP2A, UBE2C or ASPM, which are known to be oncogenic and are associated with poor patient prognosis. We also generated SRSF3 siRNA-encapsulating lipid nanoparticles as a future therapeutic tool. This study suggests a therapeutic strategy for NSCLC irrespective of the mutation status of *TP53* and TK-encoding genes.

**Summary:** Knockdown of a splicing factor SRSF3 increases cellular senescence in NSCLC cells including those with no targetable mutation of tyrosine kinases and thus may represent a novel therapeutic strategy for a hard-to-treat group of NSCLC.

## Introduction

Lung cancer is the most lethal cancer type worldwide, accounting for 25% of cancer-related deaths (1). Non-small cell lung cancer (NSCLC) constitutes approximately 80% of all lung cancers (2). Molecular targeted therapies such as EGFR-tyrosine kinase inhibitor (TKI) and ALK-TKI lead to significantly improved overall survival in advanced NSCLC patients (3,4). However, the cases with targetable mutations or rearrangements of tyrosine kinase (TK)- encoding genes only consist of about 50% of NSCLC in the western populations and about 70% in the Asian populations (5). The prognosis of advanced NSCLC patients ineligible for these targeted therapies remains poor (6). Mutations of the *TP53* gene, in particular those encoding gain-of-function missense mutant p53 proteins, are also associated with therapy resistance and poor prognosis in NSCLC (7,8).

SRSF3, an RNA-binding serine/arginine-rich splicing factor 3, has been reported to be an oncogenic factor that is highly expressed and associated with poor prognosis in various types of cancer (9,10). Consistently, inhibition of SRSF3 induces cellular senescence as a tumor-suppressive mechanism, as previously shown by us (11). In glioblastoma cells representing another hard-to-treat cancer type, siRNA-mediated or CRISPR/Cas9-mediated inhibition of SRSF3 has been shown to suppress their tumorigenic phenotypes (12,13). Prompted by these previous studies, we for the first time investigate whether siRNA inhibition of SRSF3 shows a tumor-suppressive activity in NSCLC cells without a targetable TK alteration and/or with *TP53* mutation.

## Materials and methods

### Cells and cell culture

Four NSCLC cell lines were purchased from American Type Culture Collection (ATCC, Manassas, VA); A549 in February 1989, NCI-H1975 (hereafter H1975) in January 2007, NCI-H596 (hereafter H596) in March 1998 and NCI-H322 (hereafter H322) in March 1998 (currently discontinued). They were maintained in RPMI 1640 medium (Corning, Glendale, AZ) supplemented with 10% fetal bovine serum (FBS) and 1% penicillin/streptomycin solution (P/S). A viral packaging cell line 293T/17 and a glioblastoma cell line U-87 MG were also purchased from ATCC in March 2007 and February 2006, respectively, and maintained in DMEM medium (Corning) supplemented with 10% FBS and 1% P/S. Upon receipt of these cell lines, the certificates of analysis were available from ATCC to authenticate adherent growth property, no mycoplasma contamination, no bacterial and fungi contamination, and short tandem repeat profiles. Although these cell lines were obtained years ago, the cumulative periods in cell culture in our laboratory were less than 2 months. All cell cultures were maintained in a humidified incubator with 5% CO_2_ at 37 C° and periodically confirmed to be mycoplasma-free. The mutation statuses of *TP53* and TK-encoding genes in the NSCLC cell lines are summarized in Table 1.

**Table 1.**
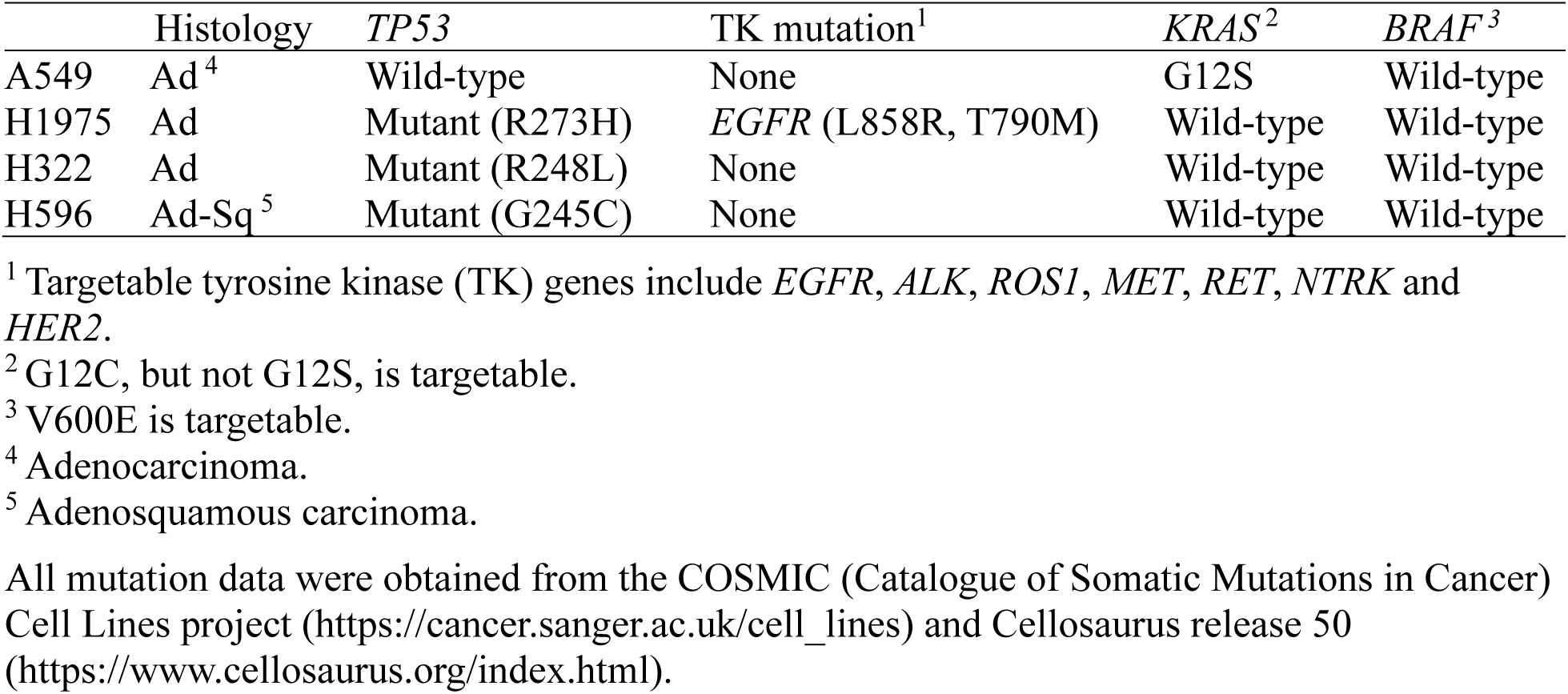
NSCLC cell lines used in this study.

### Transfection of siRNA oligonucleotides

siRNA oligonucleotides were transfected at a final concentration of 10 nM (in A549, H1975 and H322 cells) or 20 nM (in H596 cells) using Lipofectamine RNAiMAX Transfection Reagent (Thermo Fisher Scientific, no. 13778150, Waltham, MA) according to the manufacture’s instructions. Silencer Select siRNA targeting SRSF3 (#1: Assay ID s12732 and #2: Assay ID s12733) and non-targeting negative control siRNA (no. 4390843) (11) were purchased from Thermo Fisher Scientific. Cells were collected 72 h after transfection for qRT-PCR, western blot analysis, SA-β-gal staining, immunofluorescence staining, and ELISA.

### RNA isolation and qRT-PCR

Total RNA samples were prepared using RNeasy Plus Micro Kit (QIAGEN, no. 74034, Germantown MD). Reverse transcription was carried out using High-Capacity cDNA Reverse Transcription Kit (Thermo Fisher Scientific, no. 4368814). qRT-PCR assays were performed using Taqman Gene Expression Master Mix (Thermo Fisher Scientific, no. 4369016) and the following primers/probe sets (all from Thermo Fisher Scientific): SRSF3 (Hs00751507_s1); BAX (Hs00180269_m1); BBC3 (PUMA, Hs00248075_m1); ASPM (Hs00411505_m1); UBE2C (Hs 00964100_g1); TOP2A (Hs01032137_m1); and GAPDH (Hs02758991_g1) as an internal control. SYBR Select Master Mix (Thermo Fisher Scientific, no. 4472908) was used to detect p53β (Forward: 5′-GCGAGCACTGCCCAACA-3′ and Reverse: 5′-GAAAGCTGGTCTGGTCCTGAA-3′) and GAPDH (Forward: 5′-TGCACCACCAACTGCTTAGC-3′ and Reverse: 5′-GGCATGGACTGTGGTCATGAG-3′). Quantitative data analysis was performed using the ΔΔCt method according to the manufacturer’s instructions (https://assets.thermofisher.com/TFS-Assets/LSG/manuals/cms_042380.pdf). All qRT-PCR data were mean ± standard deviation (S.D.) from technical triplicate (n=3).

### Protein lysate preparation and western blot analysis

To extract protein lysates from cells, RIPA Lysis and Extraction Buffer containing 25mM Tris HCl pH 7.6, 150mM NaCl, 1% NP-40, 1% sodium deoxycholate, and 0.1% SDS (Thermo Fisher Scientific, no. 89900) with Halt Protease and Phosphatase inhibitor Cocktail (Thermo Fisher Scientific, no. 78440) was used. Bradford protein assay was performed to determine protein concentration using Bio-Rad Protein Assay Dye Reagent Concentrate (Bio-Rad, no. 5000006, Hercules, CA). Protein lysates were added with 4× Laemmli Sample buffer (Bio-Rad, no. 1610747), boiled at 95℃ for 5 min, separated on 10% or 10-to-20% Novex Tris-Glycine Mini Protein Gels (Thermo Fisher Scientific, XP00100BOX or XP10200BOX), and transferred to 0.45-μm PVDF transfer membranes (Thermo Fisher Scientific, no. 88518). Membranes were blocked using SuperBlock T20 (TBS) blocking buffer (Thermo Fisher Scientific, no. 37536). Incubations with primary and secondary antibodies were carried out in 1:1 mixture of 10 mM Tris-HCl (pH 7.5)/150 mM NaCl/0.05% Tween 20 (TBS-T) and SuperBlock T20 (TBS) blocking buffer. Signal detection was performed using SuperSignal West Dura chemiluminescence substrate (Thermo Fisher Scientific, no. 34076) and ChemiDoc Imaging System (Bio-Rad). Quantitative image analysis was performed using Image Lab software (Bio-Rad, ver. 6.1).

Primary antibodies used were as follows: SRSF3 (Santa Cruz Biotechnology, sc-13510, Santa Cruz, CA); p53 (DO-1) (Santa Cruz Biotechnology, sc-126); cleaved PARP (Asp214, Cell Signaling Technology, no. 9541, Danvers, MA); cleaved caspase-3 (Cell Signaling Technology, no. 9664); p21^WAF1^ (Cell Signaling Technology, no. 2947); p16^INK4A^ (Santa Cruz Biotechnology, sc-1661); phospho-histone H2A.X (EMD Millipore, no. 05-636, Burlington, MA); UBE2C (Santa Cruz Biotechnology, sc-271050); TOP2A (Cell Signaling Technology, no. 12286); GAPDH (EMD Millipore, MAB374); and β-actin (Abcam, ab6276, Waltham, MA). Secondary antibodies used were goat anti-mouse IgG (H+L) (Thermo Fisher Scientific no. 32430) and goat anti-rabbit IgG (H+L) (Thermo Fisher Scientific no. 32460).

### SA-β-gal staining

Senescence-associated beta-galactosidase activity (SA-β-gal) staining was performed with the Senescence Associated (SA)-β-Galactosidase Staining Kit (Cell Signaling Technology, no. 9860) according to the supplier’s protocol. Percentages of positive cells were shown as mean ± S.D. from biological triplicates, each observing at least 100 cells.

### Immunofluorescence staining

Cells were seeded in 8-chamber microscope slides (Thermo Fisher Scientific, Nunc Lab-Tek II CC2 Chamber Slide System, 8-well, no. 154739), fixed with 4% paraformaldehyde in phosphate-buffered saline (PBS), permeabilized with 0.25% Triton X-100 in PBS, blocked with 1% bovine serum albumin, and incubated at 4℃ overnight with a primary antibody: anti-p21 (Santa Cruz Biotechnology, sc-6246), anti-cleaved caspase-3 (Cell signaling Technology, no. 9664S) or anti-Ki67 (Abcam, ab56978). Incubation with a secondary antibody was for 1 h at room temperature using donkey anti-rabbit IgG (H+L) highly cross-adsorbed secondary antibody, Alexa Fluor 594 (Thermo Fisher Scientific, A-21207) or donkey anti-mouse IgG (H+L) highly cross-adsorbed secondary antibody, Alexa Fluor 488 (Thermo Fisher Scientific, A-21202). Slides were mounted with ProLong Gold Antifade Mountant with DAPI (4′,6-diamidino-2-phenylindole) (Thermo Fisher Scientific, P36931). All images were taken using a confocal microscope (Zeiss 780) with ZEN software (Zeiss, White Plains, NY).

### ELISA

Quantification of IL-6 in the cell culture media was performed using the Human IL-6 ELISA Kit (Sigma-Aldrich, RAB0306, St. Louis, MO) according to the manufacturer’s instructions. The assay was performed in a 96-well plate format, and absorbance was measured at 450 nm using a microplate reader SpectraMax ABS Plus (Molecular Devices, San Jose, CA). Standard curves were generated using recombinant human IL-6 provided in the kit, and the concentration of IL-6 in the samples was calculated accordingly.

### p53β lentiviral vectors and transduction

The FLAG-tagged p53β cDNA was previously cloned in the lentiviral vector pLOC-GFP-Blasticidin (Open Biosystems, Lafayette, CO) (14). In this lentiviral vector, a missense mutation R248Q or R273H was generated via site-directed mutagenesis using the QuikChange II XL Site-Directed Mutagenesis kit (Agilent Technologies, no. 200521, Santa Clara, CA). Lentiviral packaging was performed using Trans-Lentiviral Packaging System (Open Biosystems, TLP4614) in 293T/17 cells, followed by collection of the culture supernatant after 48 h. Lentiviral transduction to NSCLC and glioblastoma cells was carried out as previously described (14–16). Transduced cells were subjected to selection with 5-10 μg/ml blasticidin (Thermo Fisher Scientific, A1113903) at 48 h post-transduction.

### Preparation and transfection of siRNA-encapsulated lipid nanoparticles

Lipid nanoparticles (LNP) were formulated by microfluidic mixing using the lipids SM-102, distearoylphosphatidylcholine, cholesterol, and 1,2-Dimyristoyl-sn-glycero-3-methoxypolyethylene glycol 2000 at a 50:10:38.5:1.5 mol ratio. Lipids were diluted in ethanol, while siRNA was diluted in 10 mM citrate buffer at pH 4. Microfluidic mixing was performed using a NanoAssemblr^TM^ Ignite^TM^ (Cytiva) programmed to a 3:1 RNA:lipid flow rate ratio, total flow rate 20 mL/min. LNPs were diluted 1:40 in PBS + 10% sucrose, then concentrated to 1 mg/mL by centrifuge filtration. The above-mentioned SRSF3 siRNA (#1: Thermo Fisher Scientific, Assay ID s12732) or control siRNA (Thermo Fisher Scientific, no. 4390843) was encapsulated at a final concentration of approximately 10 µM. A human anti-ROR1 recombinant antibody scFv fragment (clone V353) (Creative Biolabs, Shirley, NY) was conjugated to the surface of the LNP, as previously described (17). The resulting LNP (amounts corresponding to a final siRNA concentration of 10 nM) were directly added to cell culture without a transfection reagent (18).

### Search for genes that may mediate SRSF3 knockdown-induced cellular senescence

Differentially expressed genes (DEGs) in NSCLC tissues (in relative to matched non-tumor lung tissues) were from Supplementary Table S1 in ref. (19). DEGs associated with SRSF3 inhibition in two glioma stem-like cells were from Supplementary Table S3 in ref. (13). The genes commonly present in these two groups of DEGs were extracted (Supplementary Table 1) and searched in literature for their functions in cellular senescence.

### Survival analysis

The Cancer Genome Atlas (TCGA) data of lung adenocarcinoma RNA-seq available at TCGA PanCancer Atlas (https://www.cancer.gov/ccg/research/genome-sequencing/tcga) (20) was downloaded via cBioPortal [https://www.cbioportal.org] and used for survival analysis with Kaplan-Meier method.

### Statistical analysis

All quantitative data are presented as mean ± S.D. from at least three replicates. Statistical significance was evaluated using unpaired 2-tailed Student’s *t*-test. \**P* ≤ 0.05, \*\**P* ≤ 0.01, \*\*\**P* ≤ 0.001.

## Results

### SRSF3 knockdown inhibits cell proliferation and increases cellular senescence in NSCLC cell lines

We transfected two independent siRNA oligonucleotides against SRSF3 (11), along with a control siRNA, into four NSCLC cell lines: A549 (no TK mutation; *TP53* wild-type), NCI-H1975 (*EGFR* L858R/T790M; *TP53* R273H mutant), NCI-H322 (no TK mutation; *TP53* R248L mutant) and NCI-H596 (no TK mutation; *TP53* G245C mutant) (Table 1). The siRNA-mediated knockdown of SRSF3 was confirmed in both qRT-PCR and western blot analyses (Figure 1A and B). Upon SRSF3 knockdown, all the four cell lines showed a significant increase in cells stained positive for SA-β-gal activity, a widely used marker of cellular senescence (Figure 1C). Another feature commonly observed in all the cell lines with SRSF3 knockdown was a reduction in cell proliferation, indicated by decreased Ki67-positive cells (Figure 1D) and consistent with senescent proliferation arrest. Other senescence-associated events were observed in one or more cell lines, but not all. These include: an upregulation of a p53-regulated cyclin-dependent kinase (CDK) inhibitor p21^WAF1^ in A549, H1975 and H596, but not in H322 (Figure 2A); increased secretion of a proinflammatory cytokine IL-6, representing senescence-associated secretory phenotype (SASP), in H1975 and H596, but not in the other two (Figure 2B); a remarkable accumulation of phosphorylated histone-H2A.X (γ-H2AX), an indicator of DNA double-strand breaks, in A549 but not in the other three (Figure 2C); and an upregulation of another CDK inhibitor p16^INK4A^ in A549 and H596, but not in the other two (Figure 2C).

**Figure 1.**
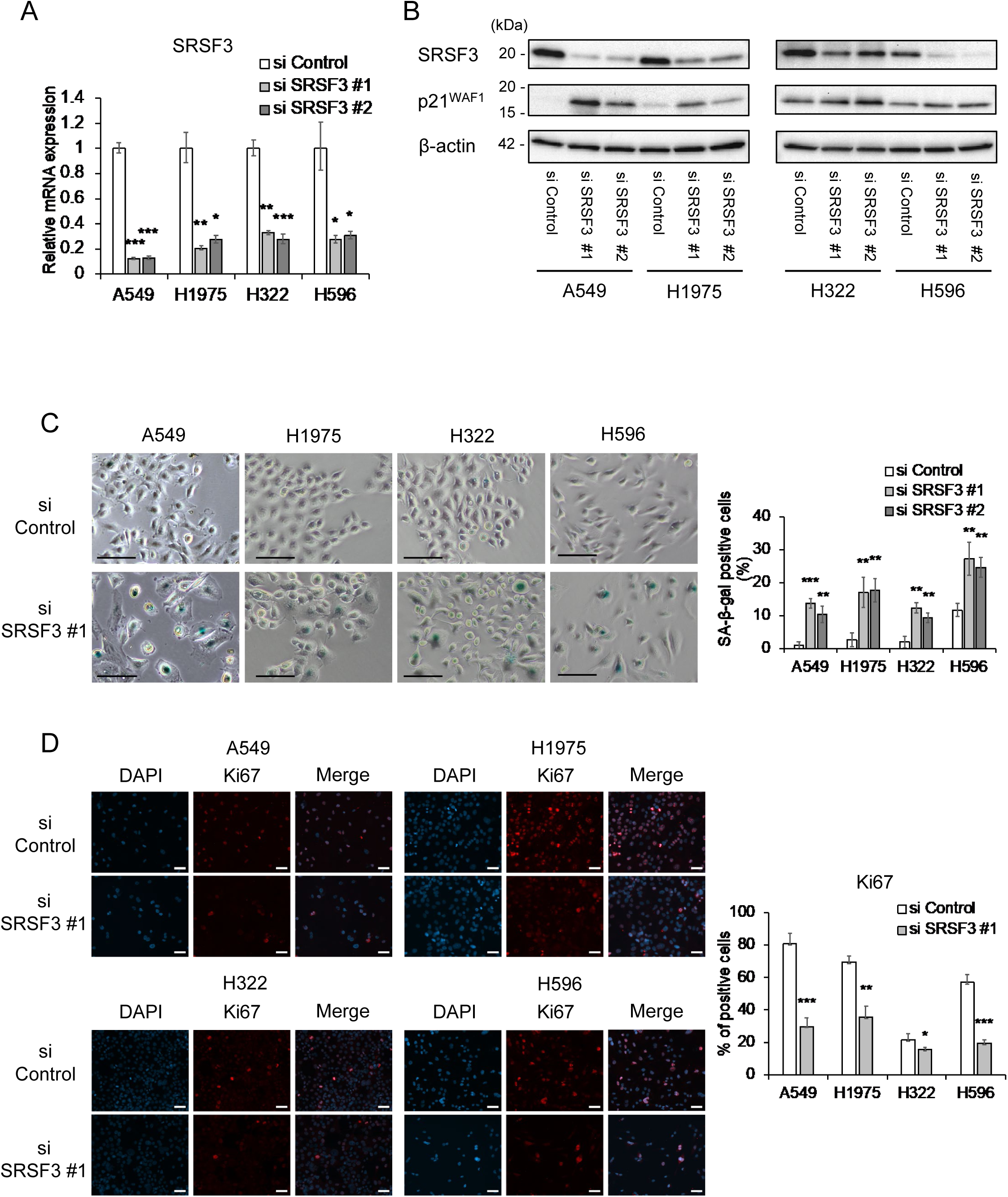
Cellular senescence induced by SRSF3 knockdown in NSCLC cells. Two siRNA oligonucleotides knocking down SRSF3 (#1 and #2), along with control siRNA, were transfected to A549, H1975, H322 and H596 cells. At 72 h post-transfection, the cells were collected or stained for the following assays. (**A**) qRT-PCR analysis of SRSF3 mRNA expression. Data were mean ± S.D. from triplicates. (**B**) Western blot analysis of SRSF3 and p21^WAF1^. β-actin was a loading control. (**C**) SA-β-gal staining. Representative images (left; scale bars, 100 µm) and quantitative summary of positive cells (right; mean ± S.D. from biological triplicates, each observing at least 100 cells). (**D**) Immunofluorescence staining of Ki67 in cells transfected with control siRNA and SRSF3 siRNA (#1). Representative images of Ki67 (red) and DAPI (blue) with their merged images (left; scale bars, 50 µm) and quantitative summary of Ki67-positive cells (right; mean ± S.D. from biological triplicates, each observing at least 100 cells). \**P* ≤ 0.05, \*\**P* ≤ 0.01, \*\*\**P* ≤ 0.001.

**Figure 2.**
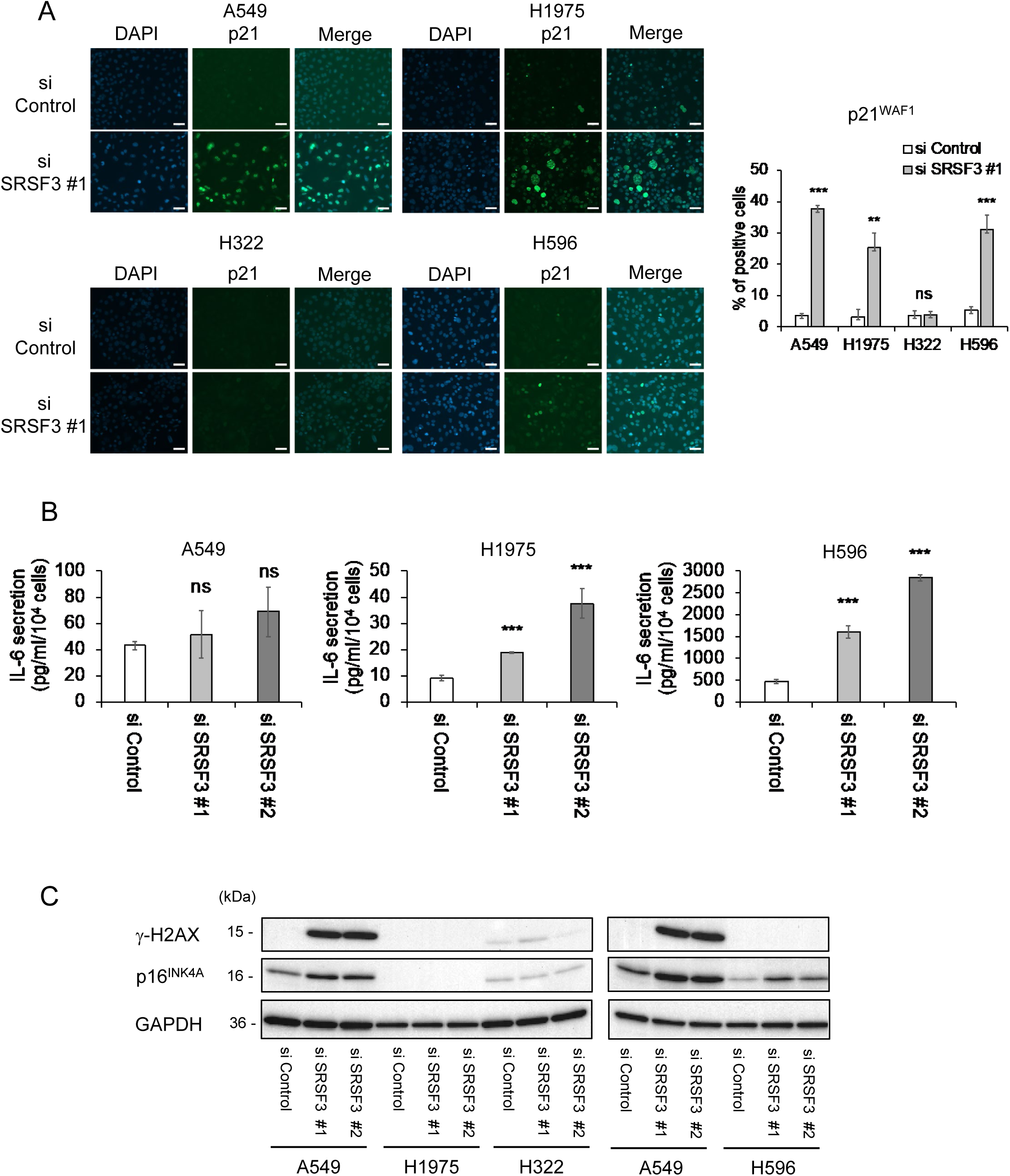
Cellular senescence-associated phenotypes in SRSF3 knocked-down NSCLC cells. (**A**) Immunofluorescence staining of p21^WAF1^ in cells transfected with control siRNA and SRSF3 siRNA (#1). Representative images of p21^WAF1^ (green) and DAPI (blue) with their merged images (left; scale bars, 50 µm) and quantitative summary of p21^WAF1^-positive cells (right; mean ± S.D. from biological triplicates, each observing at least 100 cells). (**B**) Secreted IL-6 measured by ELISA in A549, H1975 and H596 cells transfected with control siRNA and SRSF3 siRNA (#1 and #2). Data (pg/ml) were normalized to cell numbers (per 1 × 10^4^) and shown as mean ± S.D. from biological triplicates. H322 cells with either siRNA did not secrete a detectable level of IL-6. (**C**) Western blot analysis of γ-H2AX and p16^INK4A^. GAPDH was a loading control. To cross-reference two blots, three samples of A549 were included in both blots. \*\**P* ≤ 0.01, \*\*\**P* ≤ 0.001, ns = not significant.

### SRSF3 knockdown increases apoptosis in A549 cells

Since the inhibition of SRSF3 was previously reported to induce apoptosis in other types of cancer cells such as glioblastoma, colorectal cancer and ovarian cancer (12,21,22), we also examined apoptotic markers in the SRSF3 knocked-down NSCLC cell lines. The apoptotic cleavage of both caspase-3 (Figure 3A and B) and PARP (Figure 3A) was induced by SRSF3 knockdown in A549 cells, coincident with a remarkable upregulation of a p53-inducible apoptosis gene BBC3 (Figure 3C), indicating that SRSF3 knockdown in this cell line led to not only increased senescence but also increased apoptosis. Although trace levels of cleaved caspase-3 and/or cleaved PARP were detected in the other 3 cell lines (Figure 3A) and another p53-inducible apoptosis gene BAX was upregulated in H596 cells (Figure 3C), no significant increase in immunofluorescence-positive cells for cleaved caspase-3 (Figure 3B) suggests that apoptosis was not a major outcome of SRSF3 knockdown in these 3 cell lines, at least not in a cell population readily recognizable.

**Figure 3.**
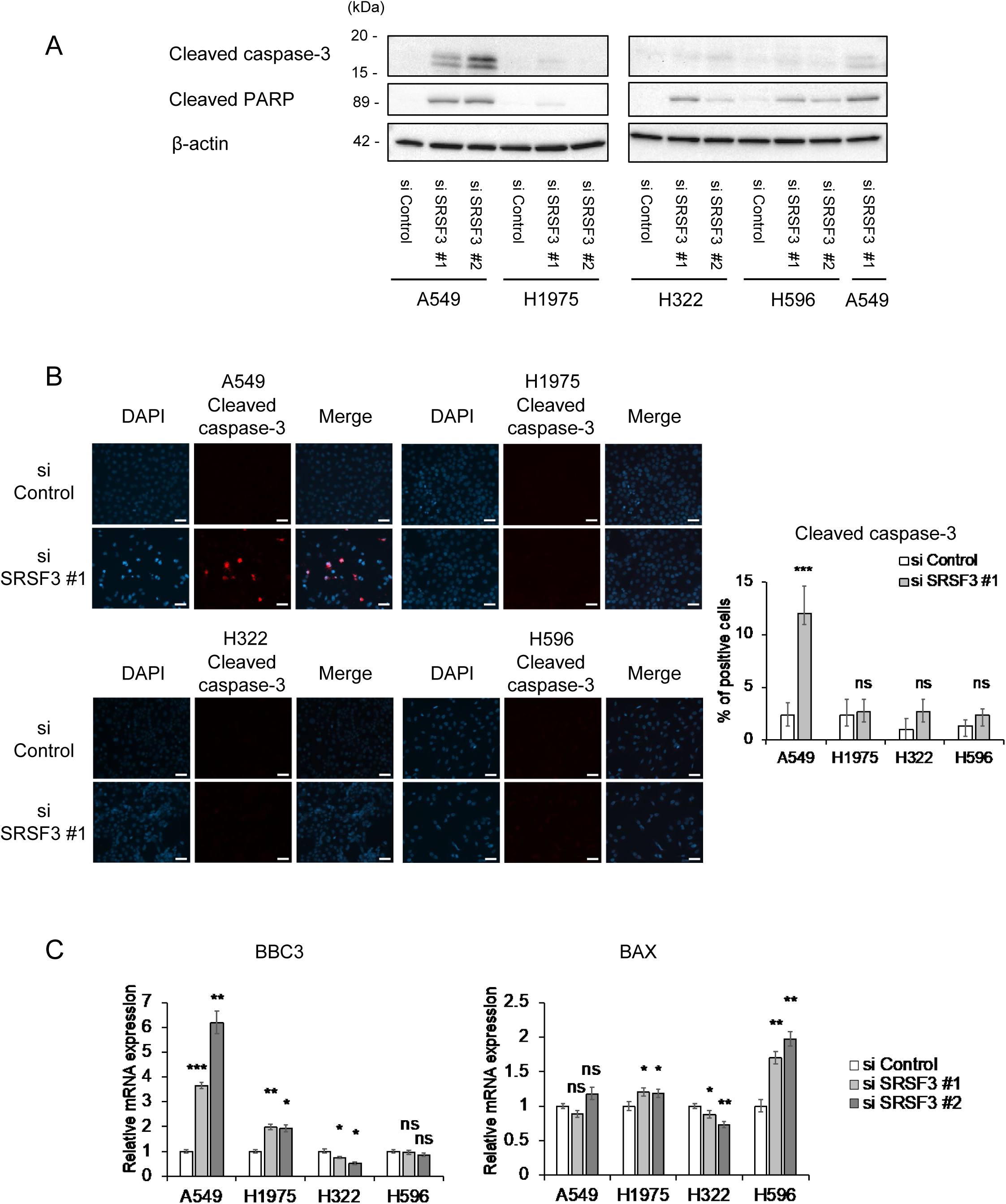
Apoptotic protein cleavage induced by SRSF3 knockdown in A549 cells. (**A**) Western blot analysis of cleaved caspase-3 and cleaved PARP in cells transfected with control siRNA and SRSF3 siRNA (#1 and #2). β-actin was a loading control. To cross-reference two blots, one sample (A549 siSRSF3 #1) was included in both blots. (**B**) Immunofluorescence staining of cleaved caspase-3. Representative images of cleaved caspase-3 (red) and DAPI (blue) with their merged images (left; scale bars, 50 µm) and quantitative summary of cleaved caspase-3-positive cells (right; mean ± S.D. from biological triplicates, each observing at least 100 cells). (**C**) qRT-PCR analysis of BBC3 and BAX mRNA expression. Data were mean ± S.D. from triplicates. \**P* ≤ 0.05, \*\**P* ≤ 0.01, \*\*\**P* ≤ 0.001, ns = not significant.

### p53β is upregulated by SRSF3 knockdown but p53β overexpression does not induce cellular senescence in NSCLC cells

In our previous reports (11,15), SRSF3 knockdown in normal human fibroblasts upregulated p53β, a natural p53 protein isoform that is generated via a SRSF3-controlled alternative mRNA splicing, which in turn induced cellular senescence in a manner dependent on wild-type p53. As previously observed in normal human fibroblasts (11), the siRNA knockdown of SRSF3 upregulated p53β in all the 4 NSCLC cell lines (Figure 4A). To directly test whether the upregulated p53β in these NSCLC cells contributes to the SRSF3 knockdown-induced senescence, we transduced A549, H1975 and H322 cells, along with U-87 MG glioblastoma cells as a control, with a constitutive lentiviral vector that drives the overexpression of p53β, which corresponds to the *TP53* status of each transduced cell line, i.e., wild-type p53β for A549 and U-87 MG, and mutant p53β at codon 273 or 248 for H1975 or H322, respectively (Figure 4B). In contrast to a significant increase in SA-β-gal-positive cells in p53β-overexpressing U-87 MG cells (N. von Muhlinen *et al*., in preparation), the overexpression of p53β in these NSCLC cell lines resulted in no increase in SA-β-gal-positive cells (Figure 4C). The apoptotic cleavage of caspase-3 and PARP, which was induced by SRSF3 knockdown in A549 cells (Figure 3A), did not occur upon p53β overexpression in any of the 3 cell lines tested, including A549 (Figure 4D).

**Figure 4.**
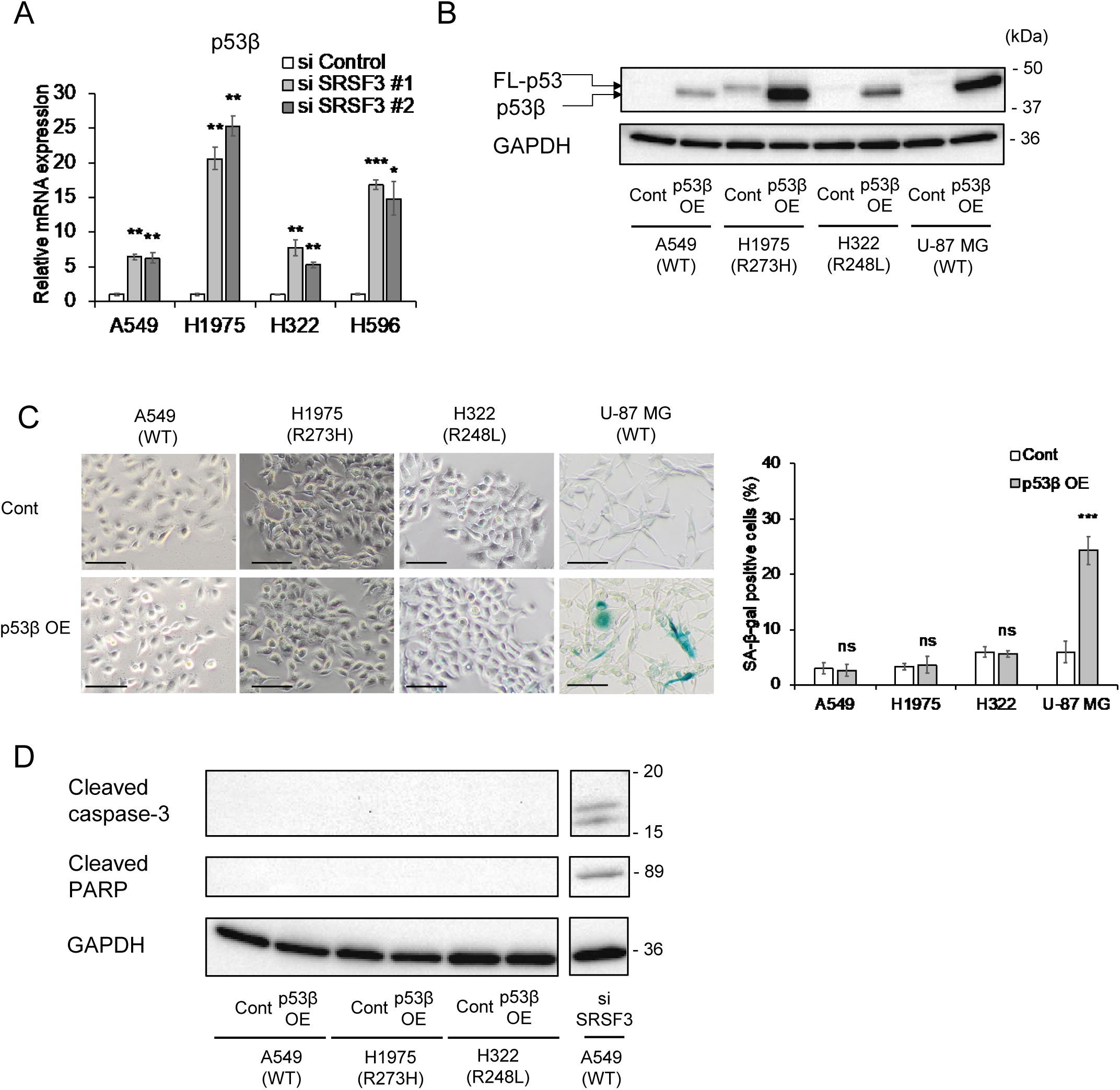
No induction of cellular senescence or apoptosis by p53β overexpression in NSCLC cells. (**A**) qRT-PCR analysis of endogenous p53β mRNA expression in NSCLC cells transfected with control siRNA and SRSF3 siRNA (#1 and #2). Data were mean ± S.D. from triplicates. (**B-D**) A549, H1975, H322 and U-87 MG (glioblastoma) cells were transduced with a lentiviral vector constitutively overexpressing p53β (p53β OE) or a control vector (Cont). The sequence of overexpressed p53β corresponded to the *TP53* status of each transduced cell line (wild-type p53β for A549 and U-87 MG, and mutant p53β at codon 273 or 248 for H1975 or H322, respectively). (**B**) Western blot analysis using anti-p53 antibody DO-1. Overexpressed p53β in all 4 cell lines and mutant full-length p53 in H1975 and H322 (FL-p53) are indicated. GAPDH was a loading control. (**C**) SA-β-gal staining. Representative images (left; scale bars, 100 µm) and quantitative summary of positive cells (right; mean ± S.D. from biological triplicates, each observing at least 100 cells). (**D**) Western blot analysis of cleaved caspase-3 and cleaved PARP in A549, H1975 and H322 cells with or without p53β overexpression. GAPDH was a loading control. A549 transfected with SRSF3 siRNA (#1 used in Figure 3A) was included in the same blot as a positive control (see the original full blot in Supplementary data). \**P* ≤ 0.05, \*\**P* ≤ 0.01, \*\*\**P* ≤ 0.001, ns = not significant.

### Repression of TOP2A, UBE2C and ASPM is associated with SRSF3 knockdown-induced cellular senescence

To identify candidate factors that may mediate the SRSF3 knockdown-induced senescence in NSCLC cells, we first compared two RNA-sequence data previously published: differentially expressed genes (DEGs) in NSCLC tissues versus matched non-tumor lung tissues (19); and DEGs associated with SRSF3 inhibition in two glioma stem-like cells (13). Five genes upregulated in NSCLC and downregulated with SRSF3 inhibition (possible oncogenes), and 6 genes downregulated in NSCLC and upregulated with SRSF3 inhibition (possible tumor suppressors) were found as candidates (Supplementary Table 1). Our subsequent literature search suggested that, among them, TOP2A (DNA topoisomerase II alpha), UBE2C (ubiquitin conjugating enzyme E2 C) and ASPM (assembly factor for spindle microtubules) were reported to play an inhibitory role in cellular senescence and a promoting role in lung carcinogenesis (23–28), which was further supported by the association between the high levels of expression of these genes and poor patient prognosis (Supplementary Figure 1). The downregulation of these 3 genes in the SRSF3 knocked-down cells was confirmed by qRT-PCR (Figure 5A: TOP2A and UBE2C in A549, H322 and H596; ASPM in all 4 cell lines) and/or western blot assays (Figure 5B: TOP2A in A549, H322 and H596; UBE2C in all 4 cell lines).

**Figure 5.**
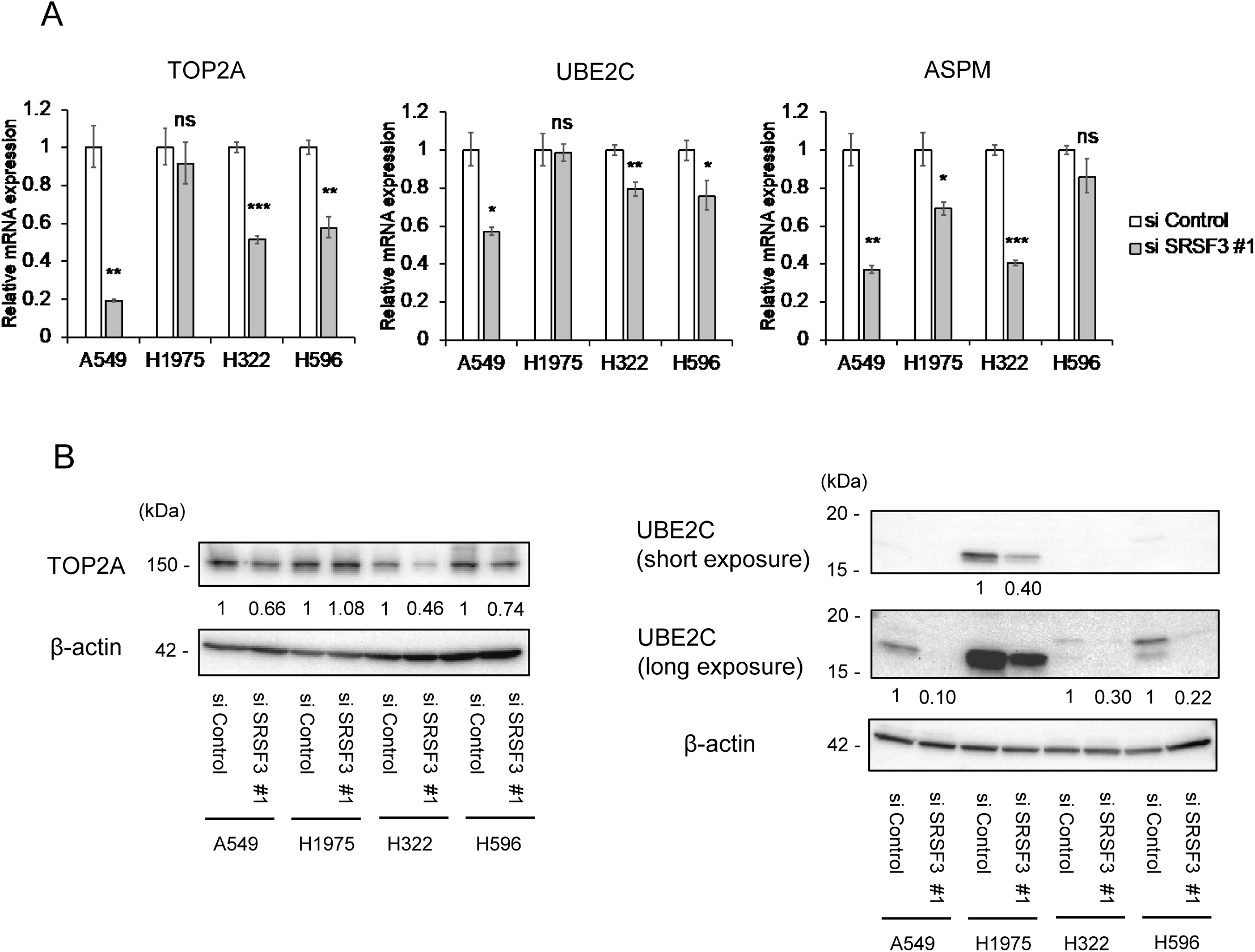
Downregulation of lung cancer-associated genes that mediates SRSF3 knockdown-induced cellular senescence in NSCLC cells. (**A**) qRT-PCR analysis of TOP2A, UBE2C and ASPM mRNA expression in NSCLC cells transfected with control siRNA (siControl) and SRSF3 siRNA (siSRSF3 #1). Data were mean ± S.D. from triplicates. (**B**) Western blot analysis of TOP2A (left) and UBE2C (right; short and long exposure images of the same blot), whose expression levels were normalized to β-actin. The relative expression values of siSRSF3 #1 to those of siControl (defined as 1 in each cell line) are shown below the images. \**P* ≤ 0.05, \*\**P* ≤ 0.01, \*\*\**P* ≤ 0.001, ns = not significant.

### Generation of SRSF3 siRNA-encapsulating lipid nanoparticles (LNP) as a future therapeutic tool

Toward future therapeutic application of the siRNA-mediated knockdown of SRSF3, we generated the LNP encapsulating SRSF3 siRNA. A simple incubation of A549 cells with the siRNA-LNP, in the absence of a transfection reagent, worked as efficiently as the standard siRNA transfection method in knocking down SRSF3 (Figure 6A and 6B), increasing p21^WAF1^ expression (Figure 6B), and inducing cellular senescence (Figure 6C), which warrants the use of this LNP in further *in vivo* studies.

**Figure 6.**
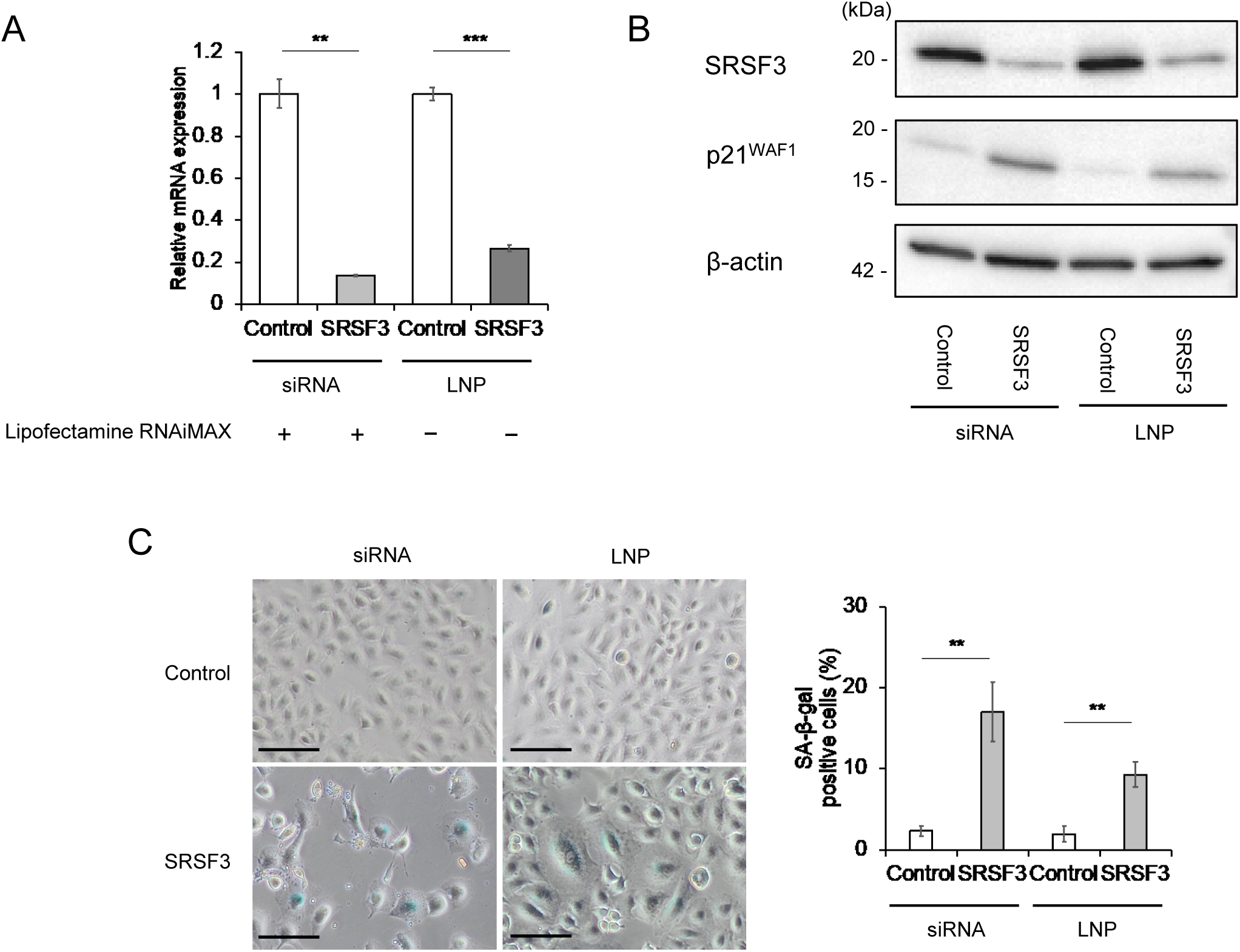
Lipid nanoparticles (LNP)-mediated activity of SRSF3 siRNA. The LNP encapsulating SRSF3 siRNA (#1) or control siRNA (at the final concentration of 10 nM as siRNA) were incubated with A549 cells in the absence of a transfection reagent (Lipofectamine RNAiMAX -), in parallel to standard transfection of SRSF3 siRNA (#1) or control siRNA (at the final concentration of 10 nM, Lipofectamine RNAiMAX +). At 72-h incubation or post-transfection, the cells were collected (A and B) or stained (C). (**A**) qRT-PCR analysis of SRSF3 mRNA. Data were mean ± S.D. from triplicates. (**B**) Western blot analysis of SRSF3 and p21^WAF1^. β-actin was a loading control. (**C**) SA-β-gal staining. Representative images (left; scale bars, 100 µm) and quantitative summary of positive cells (right; mean ± S.D. from biological triplicates, each observing at least 100 cells). \*\**P* ≤ 0.01, \*\*\**P* ≤ 0.001.

## Discussion

Our data in this study suggests that the siRNA-mediated inhibition of SRSF3 may be developed as a therapeutic approach for NSCLC with diverse genetic contexts. A strength of this study lies in the use of NSCLC cell lines with or without *TP53* mutations, with or without a mutation of TK-encoding genes such as *EGFR*, and with different histological subtypes of the original tumors (Table 1). Likely because of these variations, we observed some cell line-dependent differences in responses to siRNA knockdown of SRSF3. Increased apoptosis, indicated by cleaved caspase-3, cleaved PARP and upregulated BBC3, was observed only in A549 (Figure 3), which may be attributed to the wild-type *TP53* status (Table 1), although a caspase-3-independent apoptotic cleavage of PARP (29–31) cannot completely be ruled out in H322 and H596 (Figure 3A). Despite the line-to-line response differences in p21^WAF1^ expression (Figure 1B and 2A), IL-6 secretion (Figure 2B), γ-H2AX levels (Figure 2C) and p16^INK4A^ expression (Figure 2C), all 4 NSCLC cell lines showed an increase in cellular senescence upon siRNA knockdown of SRSF3, which was supported by both increased SA-β-gal-positive cells (Figure 1C) and reduced cell proliferation (reduced Ki67-positive cells, Figure 1D). These data suggest that the induction of cellular senescence may be a common outcome of the SRSF3 knockdown in NSCLC cells, which can be induced through one or more pathways involving the CDK inhibitors p21^WAF1^ and/or p16^INK4A^, DNA damage response, and pro-inflammatory SASP cytokines (32).

We previously reported that siRNA knockdown of SRSF3 in normal human fibroblasts increased an alternative mRNA splicing that produces p53β, a C-terminally truncated natural protein isoform of p53 (11), which in turn induced cellular senescence in these cells (11,15). In this study, p53β was shown to be induced upon SRSF3 knockdown in the NSCLC cell lines as well (Figure 4A). However, in contrast to the induction of cellular senescence by overexpression of p53β in normal human fibroblasts (15), human CD8^+^ T lymphocytes (14) and a glioblastoma cell line U-87 MG (Figure 4C; and N. von Muhlinen *et al*., in preparation), no increase in senescent cells was observed in any of the NSCLC cell lines overexpressing p53β (Figure 4C), suggesting that p53β is not a major effector of SRSF3 knockdown in NSCLC cells. Our analysis of public datasets and gene expression assays (Figure 5 and Supplementary Table 1) suggest that SRSF3 knockdown-induced senescence in NSCLC cells may be mediated by downregulation of TOP2A, ASPM and UBE2C, which all are known to act oncogenic in lung carcinogenesis (19,24,25) and whose high expression levels are significantly associated with poor prognosis (Supplementary Figure 1).

This study suggests the SRSF3 siRNA as a new therapeutic molecule for NSCLC, which may work irrespective of the mutation status of *TP53* and TK-encoding genes. However, this siRNA could also exert the same activity in normal cell types, as shown in our previous study with normal human fibroblasts (11), raising a safety concern due to normal cell senescence. To address this concern, we have designed a NSCLC-targeting LNP carrying the SRSF3 siRNA, which was shown to efficiently knock down SRSF3 and induce senescence in A549 cells (Figure 6) and will be an important material in further *in vivo* studies toward future therapeutic applications.

## Supplementary data

Supplementary data are available at *Carcinogenesis* online.

## Funding

This work was supported by Intramural Research Program of Center for Cancer Research, National Cancer Institute, National Institutes of Health. S.N. was supported by a fellowship from Nippon Medical School, Tokyo, Japan. L.Y. is supported by the JSPS Research Fellowship for Japanese Biomedical and Behavioral Researchers at NIH.

## Acknowledgements

We thank CCR Genomics Core Facility for Sanger sequencing of the lentiviral vectors and quality check of the RNA samples, and CCR Microscope Core Facility for immunofluorescence imaging. We also thank Dr. Jeffrey Cohen, National Institute of Allergy and Infectious Diseases, for helpful discussion, and Drs. Masahiro Seike and Akihiko Gemma, Nippon Medical School, Tokyo, Japan, for continuous encouragement. Graphical Abstract was created in BioRender.com.

## Author contributions

S.N. conducted the experiments, analyzed the data and drafted the paper. N.V.M. and L.Y. conducted the experiments and generated the experimental materials. J.R.M. and T.E.P. generated the LNP. H.P. and D.W. supervised the design and generation of LNP. I.H. designed the study, conducted the experiments, analyzed the data and wrote the paper. C.C.H. designed the study and supervised the overall project.

## Ethics statement

n/a

## Conflict of interest

The authors declare no potential conflicts of interest.

## Data availability

Original data in our study are available upon request.

## Abbreviations

ASPM: assembly factor for spindle microtubules
BBC3: BCL2 binding component 3
CDK: cyclin-dependent kinase
DEGs: differentially expressed genes
Dox: doxycycline
EGFR: epidermal growth factor receptor
γ-H2AX: phospho-histone-H2A.X
LNP: lipid nanoparticles
MOI: multiplicity of infection
NSCLC: non-small cell lung cancer
PARP: poly(ADP-ribose) polymerase
qRT-PCR: quantitative reverse transcription-PCR
SA-β-gal: senescence-associated β-galactosidase activity
SASP: senescence-associated secretory phenotype
siRNA: small interfering RNA
SRSF3: serine and arginine rich splicing factor 3
TK: tyrosine kinase
TKI: tyrosine kinase inhibitor
TOP2A: DNA topoisomerase II alpha
UBE2C: ubiquitin conjugating enzyme E2 C

---

**Supplementary Table 1.** Differentially expressed genes (DEGs) commonly reported in SRSF3 inhibition and in NSCLC development.

| Down in SRSF3 inhibition<sup>1</sup><br>Up in NSCLC<sup>2</sup> | Up in SRSF3 inhibition<sup>1</sup><br>Down in NSCLC<sup>2</sup> |
| --- | --- |
| ASPM | MFAP4 |
| UBE2C | IGSF10 |
| RRM2 | DNAAF1 |
| TOP2A | AOC3 |
| IGFBP3 | FOSB |
|  | ABI3BP |

<sup>1</sup> Downregulated or upregulated DEGs in SRSF3 inhibition were from Song *et al.* (ref. (13), Supplementary Table S3), which compared wild-type and SRSF3-knocked-out glioma stem-like cells.
<sup>2</sup> Upregulated or downregulated DEGs in NSCLC development were from Wang *et al.* (ref. (19), Supplementary Table S1), which compared NSCLC tissues with matched non-tumor tissues.

**Supplementary Figure 1.**
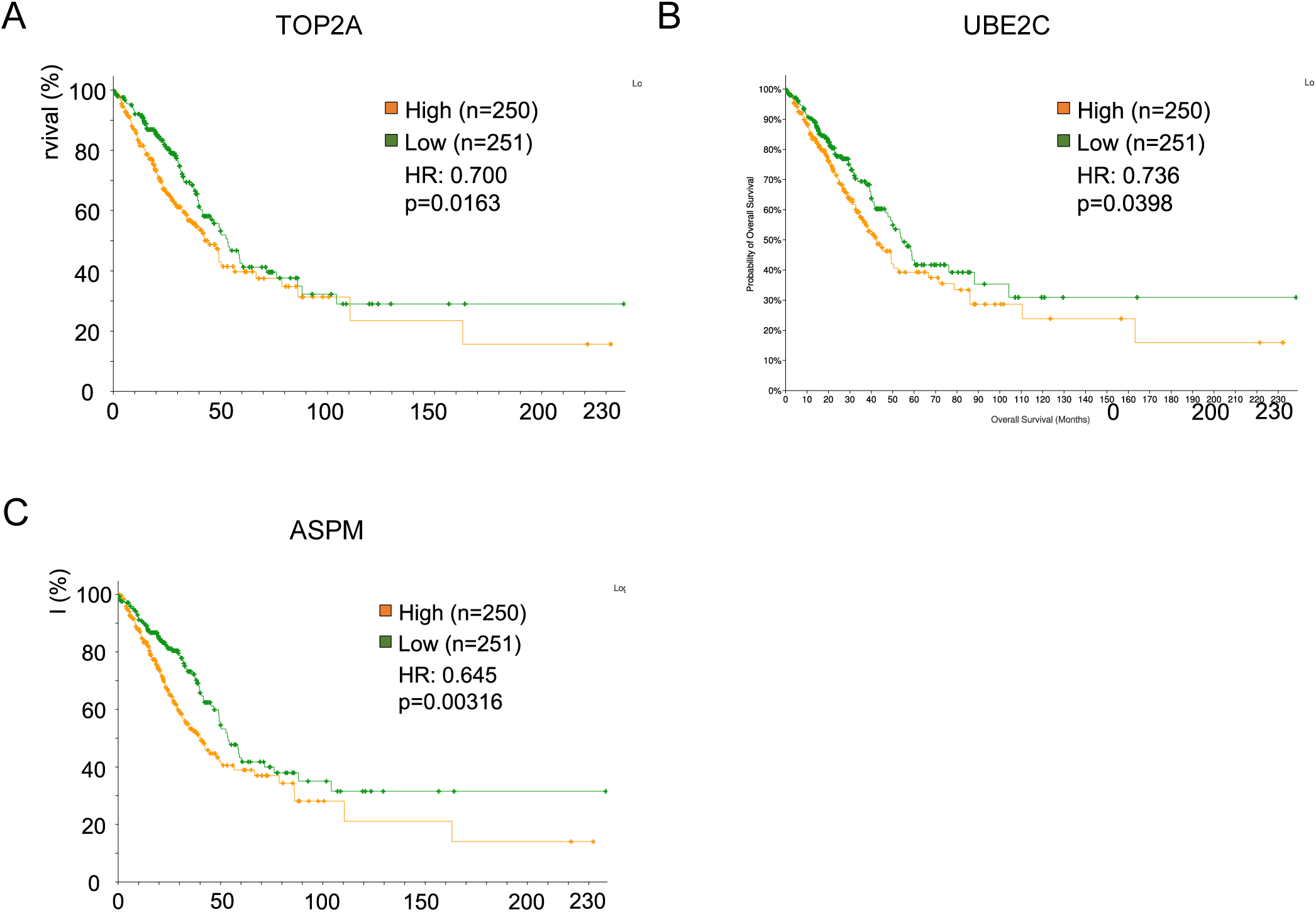
Poor patient prognosis associated with high levels of expression of TOP2A (**A**), UBE2C (**B**) and ASPM (**C**). The TCGA data of lung adenocarcinoma RNA-sequencing were analyzed for overall survival in high-level group (n=250) versus low-level group (n=251) with the Kaplan-Meier method. HR, hazard ratio.

## Notes

### Competing Interest Statement

The authors have declared no competing interest.

